# Inhibition of influenza virus replication by artificial proteins (αReps) targeting its RNA-polymerase

**DOI:** 10.1101/2024.07.03.601934

**Authors:** Mélissa Bessonne, Jessica Morel, Quentin Nevers, Agathe Urvoas, Marie Valerio-Lepiniec, Thibaut Crépin, Philippe Minard, Bernard Delmas

## Abstract

Seasonal epidemics and pandemics caused by influenza A viruses still represent a main public health burden in the world. Influenza viruses replicate and transcribe their genome in the nucleus of the infected cells, two functions that are supported by the viral RNA-dependent RNA-polymerase (FluPol) through extensive structural rearrangements and differential interactions with host cell factors. To get insights into its functioning, we screened a phage-display library of biosynthetic proteins (named αReps and build on a rigid alpha-helicoidal HEAT-like scaffold) against the structurally invariant FluPol core and several flexibly-linked domains of the FluPol PB2 subunit. Several αReps specific of the cap binding domain [CBD], the 627-domain and the NLS domain of the PB2 FluPol subunit displayed FluPol inhibitory and virus neutralization activities when transiently expressed in the cytosol. Furthermore, intracellular ectopic inducible expression of the αReps C3 and F3 (specific of the CBD and the 627-domain, respectively) in influenza virus permissive cells blocked transcription and multiplication of viruses representative of the H1N1, H3N2 and H7N1 subtypes, even when induced at late times post-infection. A synergic inhibitory effect on FluPol activity and virus multiplication was evidenced when the two αReps were covalently linked. These results suggest that i) interfering with FluPol structural rearrangements that are concomitant to its various activities may represent a promising strategy to block virus multiplication and to design new types of antivirals such as dual binders targeting distant sites on FluPol and ii) the 627-domain could be efficiently targeted to design influenza antivirals.

**Author Summary:** The influenza virus RNA-polymerase (FluPol) ensures genome transcription and replication in the nucleus of the infected cells. To select ligands able to interfere with FluPol functions, we screened a library of phages encoding biosynthetic proteins (named αReps) for binding to FluPol subunits and domains. When expressed intracellularly, several of them display efficient FluPol blocking and virus neutralizing activities. αReps C3 and F3 assembled through covalent linkages blocked FluPol activity more efficiently than their precursors. These αReps impaired multiplication of H1N1, H3N2 and H7N1 viruses, showing that their binding sites may constitute effective targets for new antiviral development.

## Introduction

Human influenza A viruses (IAV) are important viral respiratory pathogens that cause significant morbidity and death every year in the world, particularly among high risk groups including the elderly and those with serious medical conditions. Avian and porcine influenza viruses are responsible of substantial economic losses in both poultry and meat industries, also representing a burden if they can adapt and spread in humans as observed in the last H1N1 (2009) pandemics. The genome of influenza viruses is segmented, each of the eight RNA segments is encapsidated by the viral nucleoprotein NP, forming an RNA-NP complex (RNP) that includes the RNA-dependent RNA-polymerase (FluPol) bound to its promoter, which consists of the conserved 5’ and 3’ extremities of the viral RNA (vRNA).

FluPol is made by the association of three subunits of similar sizes (the two basic subunits PB1 and PB2 and the acidic protein PA) and ensures nucleotide polymerization for both replication and transcription in the nucleus of infected cells. During the latter, an additional “cap-snatching” function is implemented to steal short 5’-capped RNA primers from host mRNAs. After binding on the cell surface, virions are endocytosed and fuse with the endosomal membrane. RNPs are then released in the cytoplasm and transported to the nucleus where they start synthesizing viral mRNAs to produce viral proteins in the cytoplasm. The polymerase subunits are then imported into the nucleus and assembled into a functional trimer. Based on *in vitro* assembly and cellular localization studies, it was shown that PA and PB1 form a dimer in the cytoplasm, which is imported into the nucleus separately from PB2 (Deng et al., 2005; Huet et al., 2010). Once in the nucleus, the PA-PB1 dimer associates with PB2 to form the heterotrimeric RNA-polymerase.

The PB1 subunit functions as the polymerase catalytic subunit. It presents the conserved motifs and finger and palm subdomains characteristic of negative-strand RNA-dependent RNA polymerases, binds to the promoter sequences of the viral RNAs, and catalyzes RNA chain elongation (Wandzik et al., 2020) and references therein). The PA subunit is divided into two structurally well-defined domains separated by a 60-amino-acid-long linker, the endonuclease (amino acids 1 to 197) and a large C-terminal domain (amino acids 257 to 716) deeply associated to the PB1 subunit (Pflug et al., 2014; Reich et al., 2014). The PA endonuclease and the PB2 cap-binding domain (CBD, amino acids 320 to 483) act synergistically to promote cap-snatching-dependent transcription. The PB2 subunit is divided into the N-terminal third (PB2-N, residues 1 to 247) and the C-terminal two-thirds (PB2-C, residues 248-760) which is constituted by independently folded domains including the CBD, which is an insertion into the mid-link domain followed by the PB2 627 and NLS domains (**Fig. 1A**; Pflug et al., 2014).

**Figure 1:**
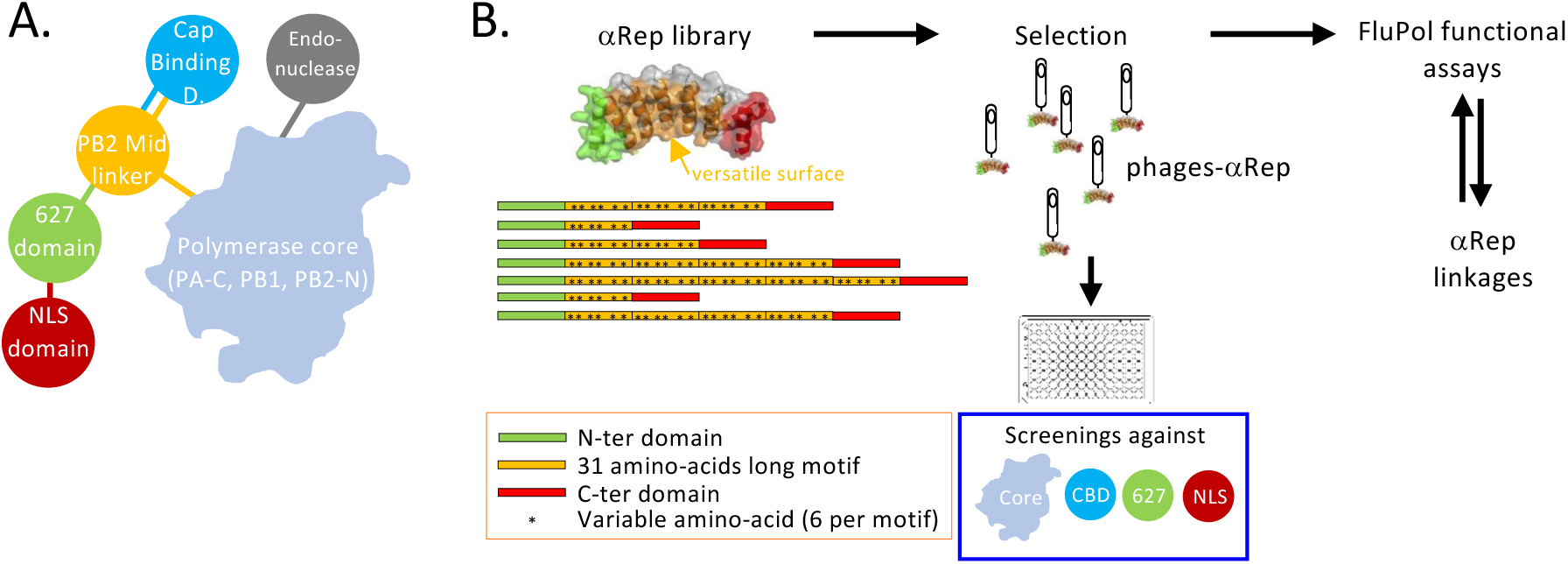
(A) Scheme showing the polymerase core and flexibly-linked peripherical domains. While the core is constituted by the PB1 subunit with PA-C and PB2-N, flexibly-linked peripherical domains are represented in circles. (**B**) Selection and characterization of anti-FluPol αReps. Screening an αReps phage library allowed the identification of several binders specific of the core and peripherical domains of FluPol. The inhibitory activity of selected intracellular αReps was evaluated using a minigenome replication assay and in virus neutralization assays.

Crystal and numerous cryo-EM structures of influenza A, B, and C RNA-polymerases and their biophysical characterization in solution have shown that FluPol has considerable conformational flexibility, and exists in several distinct functional states (Pflug et al., 2017). To summarize, these studies show that there is an invariant core comprising PB1 stabilized by the PA linker, PA-C and PB2-N, whereas the flexibly-linked peripherical domains, the PA endonuclease and the PB2-C mid-link, cap-binding, 627- and NLS domains can pack together in strikingly different configurations, depending of the binding of its promoters. The most striking example is the NLS domain, which in the c5’ promoter-bound structure has separated from 627-domain and translated about 95 Å to interact with the endonuclease domain (Thierry et al., 2016).

To gain information on FluPol functioning, nanobodies/VHH have been selected and tested for their ability to inhibit its RNA-polymerase activity (Bessonne et al., 2024; Keown et al., 2022). Discrete domains defined by neutralizing nanobodies have been mapped on the surface of FluPol. Interestingly, two nanobodies isolated from these two studies recognized an epitope that overlap the ANP32A and the RNA-polymerase II C-terminal domain binding sites on FluPol, thus blocking both replication and transcription activities and displaying a marked inhibition of virus multiplication.

As an alternative approach to VHH to identify additional vulnerability sites on FluPol, we screened a phage library encoding a family of biosynthetic proteins, named alphaReps (αReps). In contrast to VHH that display a versatile surface through their flexible CDR1-3, αReps are designed to provide a hypervariable concave rigid surface (Valerio-Lepiniec et al., 2015). αReps are constituted by alpha-helicoidal HEAT-like repeats (n x 31-amino acids long) commonly found in eukaryotes and prokaryotes including thermophiles (Andrade et al., 2001)(Urvoas et al., 2010). Sequences of homologs form a sharply contrasted sequence profile in which most positions are occupied by conserved amino acids whereas other positions appear highly variable, generating a versatile binding surface of n x 6-amino acids. A large αRep library has been assembled and was demonstrated on a wide range of unrelated protein targets to be a generic source of tight and specific binders (Valerio-Lepiniec et al., 2015). Thus, αReps were previously selected as interactors of the receptor binding domain of the spike of SARS-CoV-2 and shown to block infection of cultured cells and in hamsters (Thébault et al., 2022), or of HIV-1 nucleocapsid and to negatively interfere with virus maturation (Hadpech et al., 2017).

Here, we first obtained series of αReps specific of structurally-defined domains of FluPol, the core constituted by PB1, PA-C and PB2-N, and the flexibly-linked domains (the cap-binding domain/CBD, the 627- and the NLS domains). Several of these ligands specific of the flexibly-linked domains blocked FluPol RNA-polymerase transcriptase activity when co-expressed in cultured cells. Block of virus replication was efficient even when their expression was induced late after infection and on different virus subtypes. The covalent assembly two αReps specific of the CBD and the 627-domain resulted in a higher specific activity than their monomeric counterparts.

## Results

### Generation of αReps specific to domains of the influenza virus RNA-polymerase (FluPol)

An overview of the selection process to generate anti-FluPol αReps specific of structurally independent domains is shown in **Fig. 1B**. In order to select FluPol binders, the FluPol core (made by PB1, PA-C and PB2-N [1-116]), CBD and the 627- and NLS-domains were used as baits for screenings (**Fig. 2** for PAGE of purified FluPol polypeptides and domains and their sequences in **S1 Fig**). The phage-display procedure included three rounds of panning followed by a screening step by phage-ELISA on individual clones. Nucleotide sequencing allowed the identification of 4 independent αRep clones for the FluPol core, 9 clones for the CBD, 13 clones for the 627-domain and 17 clones for the NLS-domain that were retained for further characterizations (selected αReps and their amino acid sequences are listed in **S2 Fig**). αReps open reading frames were fused to nucleotide sequence encoding the HA-tag and placed under the control of an eukaryotic promoter to govern their expression.

**Figure 2:**
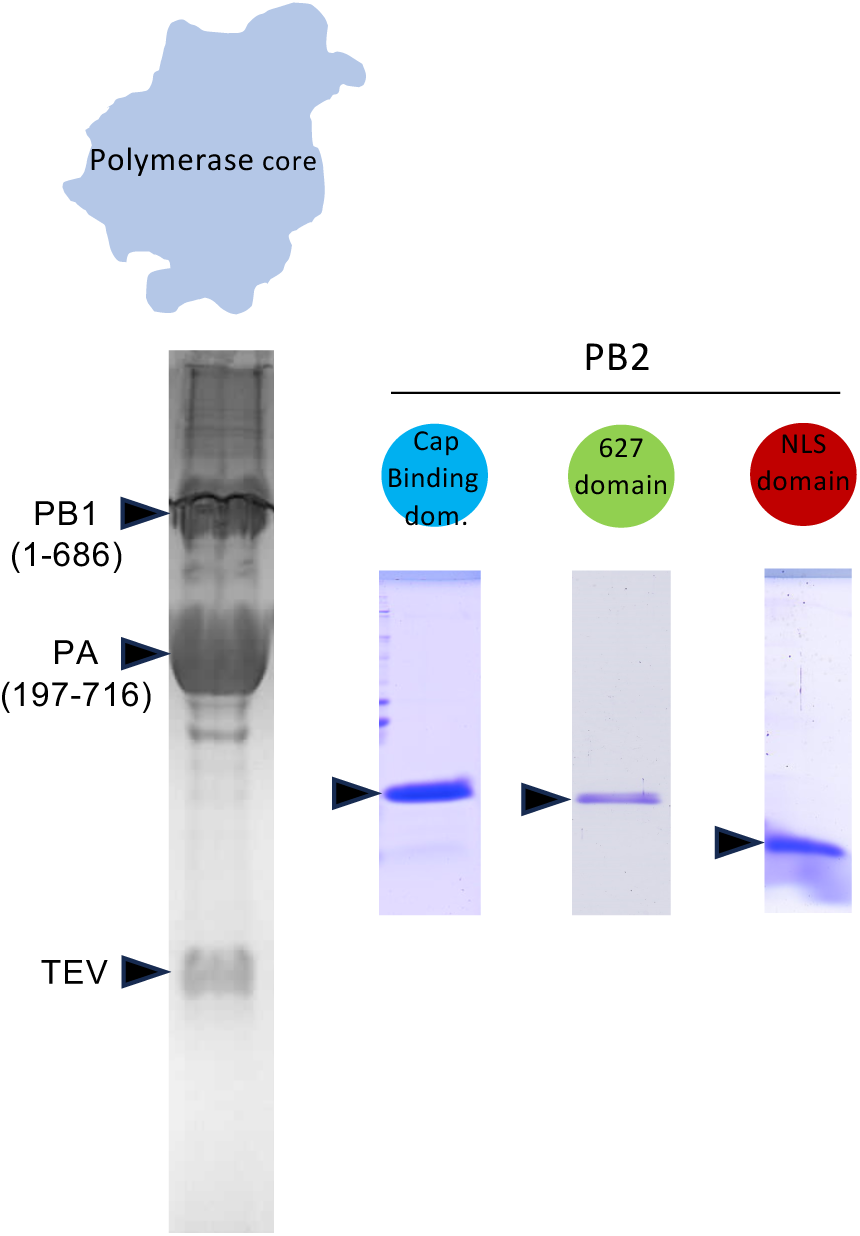
Purified FluPol subunits and domains used for αRep screenings. SDS-PAGE analysis of the purified FluPol core and of the purified PB2 domains, CBD, 627-domain and NLS domain).

### αReps specific of the flexibly-linked domains of FluPol inhibit polymerase activity *in cellulo*

To assess the inhibitory effect of αReps on polymerase activity, we used an *in cellulo* minireplicon luciferase assay in HEK-297 cells (Da Costa et al., 2015). In this assay, each of the αReps was expressed along with the polymerase subunits and NP from the influenza virus WSN strain and the pPol1-WSN-NA-firefly luciferase plasmid transcribing a modified influenza NA viral RNA (vRNA) in which the NA coding sequence is replaced by the firefly luciferase gene. The luciferase enzymatic activity measured in cell extracts reflects the overall transcription and replication activities of FluPol in the presence of an αRep expressed intracellularly. The assay revealed that several αReps specific of the CBD and the 627- and NLS-domains display potent inhibition of FluPol activity (**Fig. 3**). While only a single anti-CBD αRep (αRep C3) exhibited a significant inhibitory activity, a large proportion of anti-627 αReps displays efficient inhibition. None of the four αReps specific of the FluPol core shows an inhibitory effect (**Fig. 3A**). No anti-NLS αRep were found to display a marked inhibitory activity (**Fig. 3B**).

**Figure 3:**
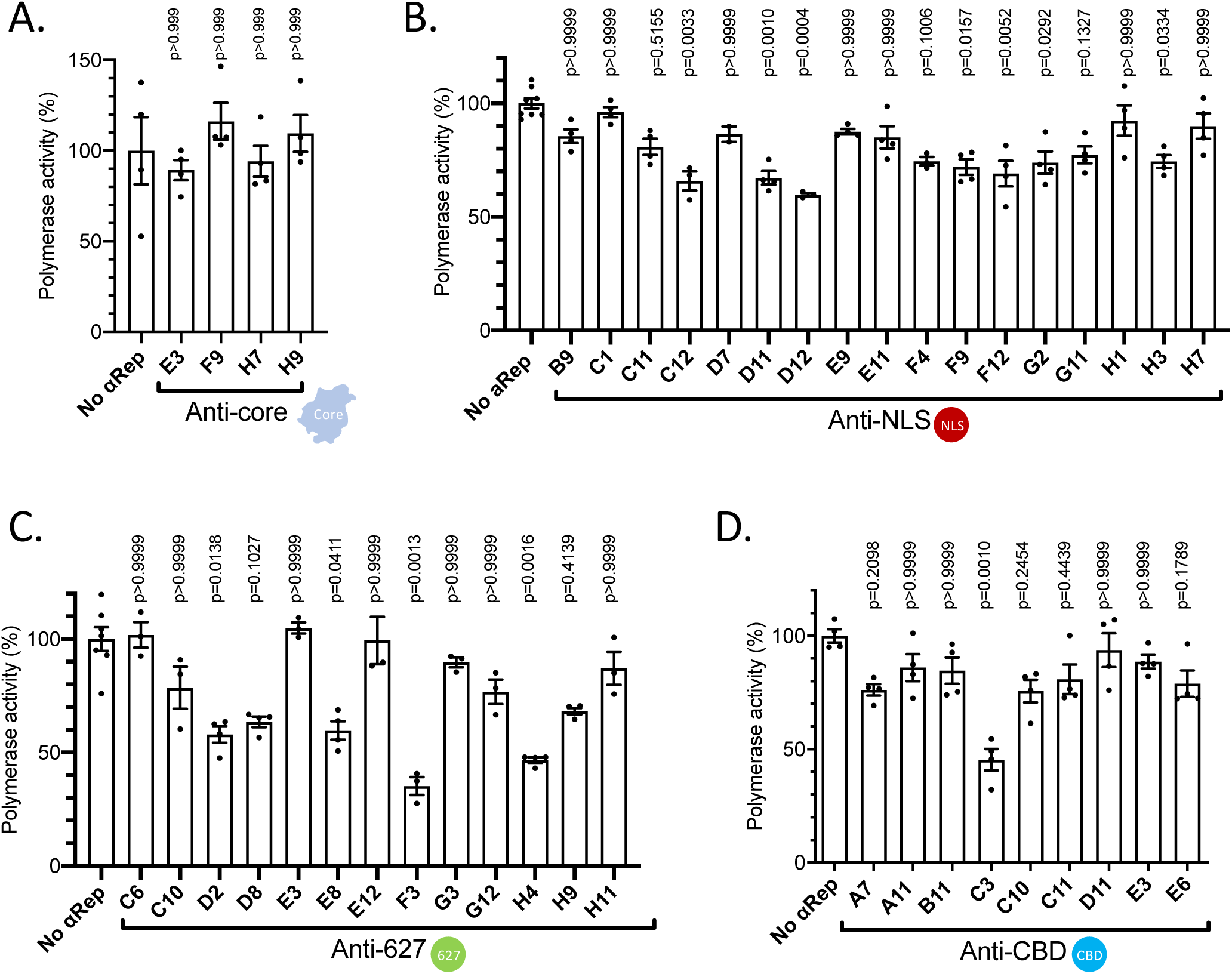
Effect of αReps expression on the FluPol replication + transcription activity. The inhibition activity of αReps on FluPol functioning was quantified in a luciferase-reporter minireplicon assay. (**A**) With anti-PA-PB1 dimer αReps; (**B**) With anti-NLS domain αReps; (**C**) With anti-627 domain αReps; (**D**) With anti-cap binding domain (CBD) αReps. Plasmids expressing NP, PA, PB1, PB2 of the WSN strain (an influenza A H1N1 virus) were co-transfected in HEK 293T cells together with the WSN-NA-firefly-luciferase reporter plasmid and a plasmid encoding an αRep. A plasmid encoding the nano-luciferase was co-transfected to control DNA uptake and normalize minireplicon activity. Luciferase activities were measured in cell lysates 48 hours post-transfection. Data are mean of ± s.e.m. n= 3 or 4 technical replicates. Kruskas-Wallis test was used to compare luminescence in the presence or absence of aReps. P<0.05 is considered significant.

Based on these data, we selected two inhibitory αReps, C3 specific of the CBD and F3 specific of the 627-domain to validate their cellular expression, their ability to recognize the full-length PB2 and to further characterize their FluPol inhibitory activities. Expression of the two αReps was validated by an indirect immune-fluorescence assay in HEK-293 cells (**S3-Fig**). Co-immunoprecipitation - western blot assays with epitope-tagged versions of PB2 and αReps confirmed that C3 and F3 are able to bind their target in the full-length form of PB2 (**Fig. 4A**). **Fig. 4B** shows a dose-response curve with C3 and F3 αReps that exhibit similar apparent specific activities on FluPol.

**Figure 4:**
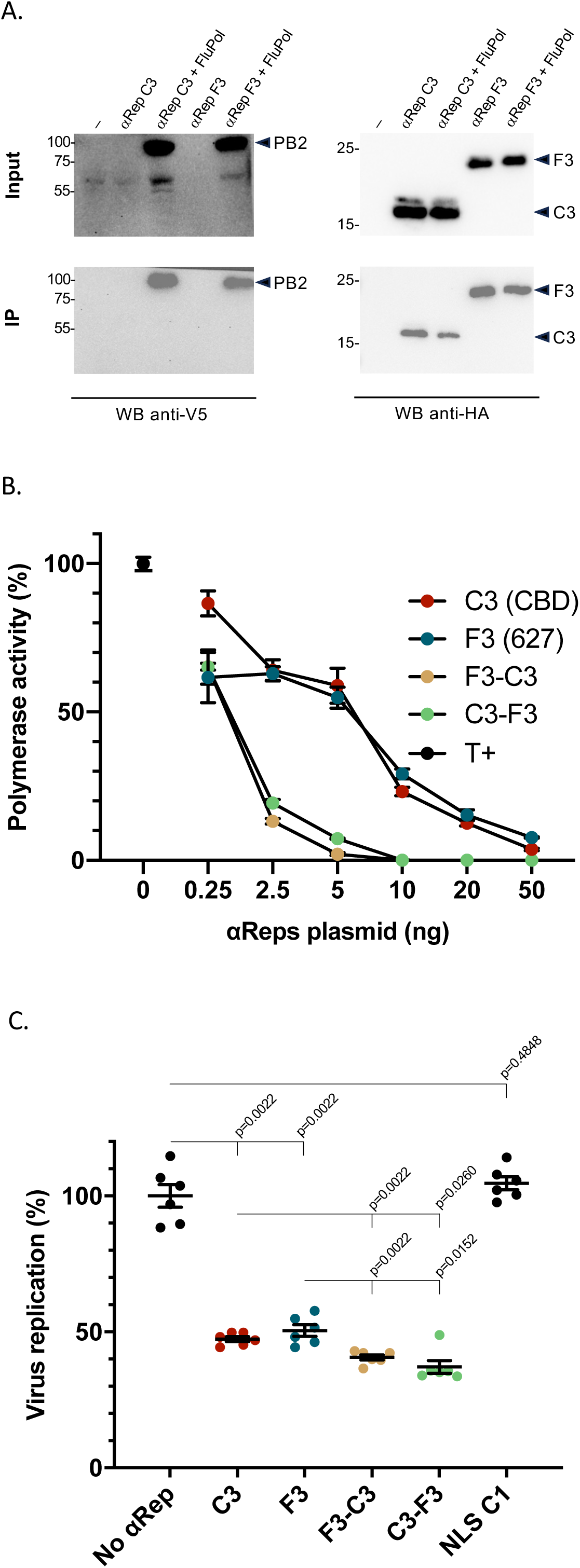
The αReps C3 (anti CBD) and F3 (anti 627-domain). **(A)** Ability of the two αReps to recognize the full-length PB2 using a coimmunoprecipitation assay. Plasmids encoding the HA-αReps and PB2-Flag were transfected in HEK 293T cells. Twenty-four hours post-transfection, cells were lysed and immunoprecipitation was carried out with an anti-HA antibody. Immunoprecipitated proteins were revealed using anti-V5- and anti-HA-tag antibodies. **(B)** The FluPol RNA-polymerase activity *in cellulo* in presence of αReps C3 and F3, and the αRep heterodimers C3-F3 and F3-C3 was monitored using a luciferase-reporter minireplicon assay as described in Fig. 2 except that the amount of transfected plasmids was normalized with the expression vector without encoded αRep. Data are mean ± s.e.m. n=2 independent transfections with n=3 technical replicates (**C**) Effect of αReps C3, F3, C3-F3 and F3-C3 on the replication of A/WSN/1933 reporter virus encoding nanoluciferase (H1N1-Nluc) (Tran et al., 2013) as a fusion PA-Nluc protein in HEK cells. An anti-NLS domain αRep (C1) was used as a negative control. Data are mean ± s.e.m. n=2 independent infections with n=3 technical replicates. Mann-Whitney test was used to compare luminescence in the presence of the αReps and derivates.

### The αReps C3 (anti-CBD) and F3 (anti-627 domain) impair infection

Next, we explored the ability of αReps C3 and F3 to inhibit or block virus multiplication. For this purpose, to quantify viral multiplication, we took advantage of a recombinant bioluminescent reporter virus with the nanoluciferase gene inserted in frame into the PA genomic segment (Tran et al., 2013). We first assessed the inhibitory effect of the αReps in a transient transfection assay in HEK-293T cells, in which αRep-encoding plasmids were transfected 24 hrs before infection. **Fig. 4C** shows that the C3 and F3 constructs were able to partially block virus multiplication.

To validate their ability to inhibit virus replication, we aimed at determining the virus susceptibility of RK13 cell lineages allowing αRep inducible expression. Plasmids in which the C3 and F3 open reading frames were fused to sequences encoding a 2A self-cleaving peptide and GFP and placed under the control of the P_TRE3G_ promoter (which is inducible by doxycycline) were constructed (**Fig. 5A**). Two RK13 cell clones were selected on their ability to express each αRep in an inducible manner (**S4 Fig**). **Fig. 5B** shows that virus multiplication was inhibited by doxycycline addition in a dose-dependent manner for both αReps, F3 being slightly more effective than C3. Next, we quantified the inhibition activity of the two αReps when expressed at early times or at late times post-infection. As shown in **Fig. 5C** and as expected, virus inhibition (measured at 24 hrs post-infection) was more effective when doxycycline was added before infection than later. However, a half-maximal inhibitory effect was observed at late times post-infection. αRep F3 expression induction at six hours post-infection still blocked 50% of virus multiplication.

**Figure 5:**
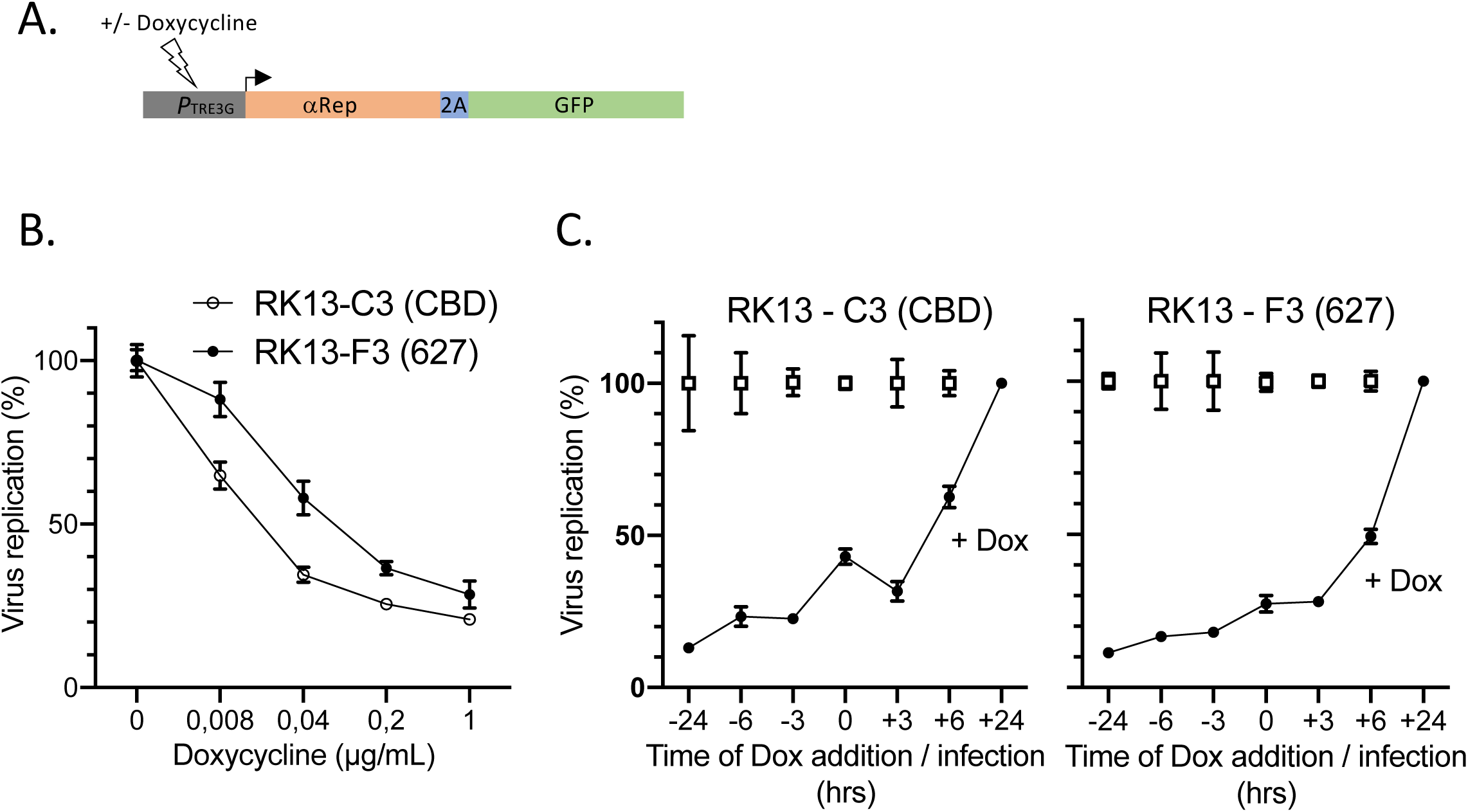
The αReps C3 (anti-CBD) and F3 (anti-627 domain) impair influenza virus replication at late time points. **(A)** Scheme of the construct allowing the inducible (+/- doxycycline) expression of the open reading frame encoding the αRep-2A-GFP polyprotein. (**B**) Quantification of the H1N1-Nluc reporter virus replication in RK13 cell clones expressing either C3 or F3 αRep in a doxycycline (Dox)-inducible manner. Data are mean± s.e.m. n=2 independent infections with n=3 technical replicates (**C**) RK13 cells expressing C3 or F3 αReps were seeded 48 h before infection with the H1N1-Nluc virus. αRep expression was induced at the indicated time points relative to infection. Cell lysates were collected 24 h p.i., and virus replication was quantified by measurement of the Nluc enzymatic activity. Mock-treated cells were included at each indicated time point relative to infection. Data are mean± s.e.m. n=1 independent infections with n=3 technical replicates.

To further investigate the activity spectrum of αReps C3 and F3, we determined their inhibitory activities towards two additional influenza viruses belonging to the H7N1 and H3N2 subtypes (A/Turkey/Italy/977/1999 [H7N1] and human influenza A/Scotland/20/1974 [H3N2]), and both tagged with the nanoluciferase (Nluc) reporter gene (**Fig. 6**). As found with the H1N1 (WSN strain) reporter virus on two independent C3 and F3 RK13 clones, multiplication of the two H7N1 and H3N2 viruses was blocked efficiently when expression was induced, suggesting sequences conservation of the C3 and F3 binding sites on their different FluPols. Here again, F3 was slightly more effective than C3. Multiplication of vesicular stomatitis virus (a virus unrelated to influenza viruses) was not altered after doxycycline induction, showing the specificity of the inhibition of influenza virus multiplication and the absence of toxicity of the induced αRep expression.

**Figure 6:**
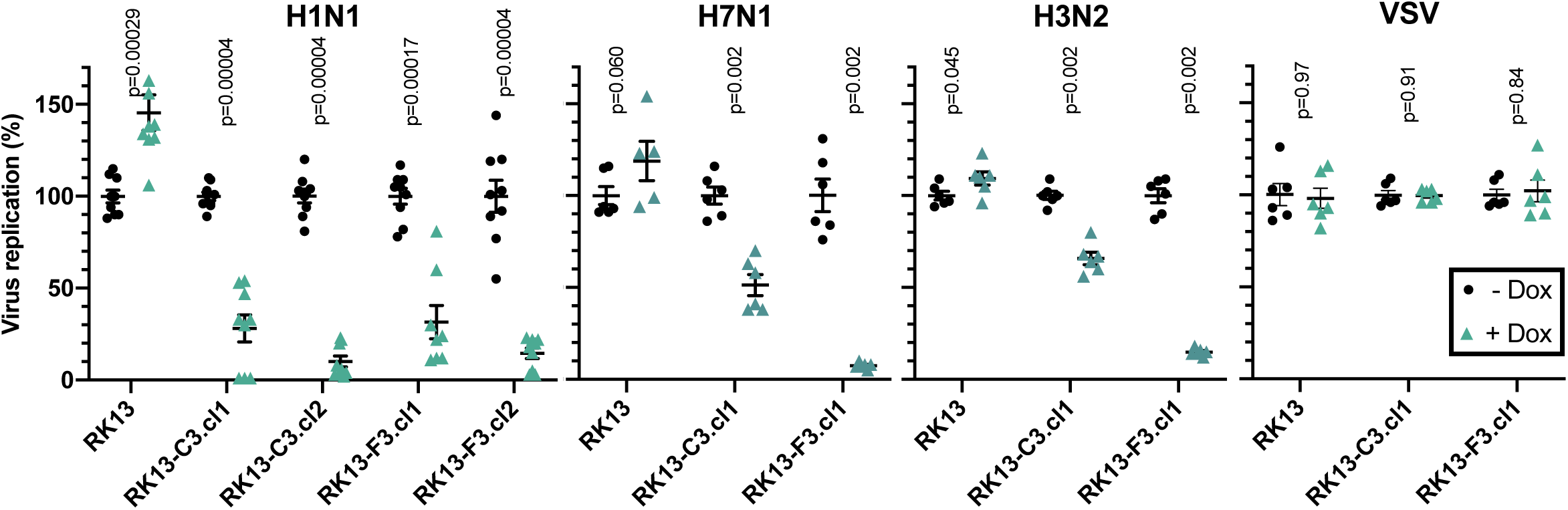
C3 and F3 αReps impair replication of viruses of H1N1, H7N1 and H3N2 subtypes. C3- and F3-derived RK13 clones (cl1 and cl2) were infected with influenza viruses of different subtypes (H1N1, H7N1 and H3N2) and encoding a reporter nanoluciferase (Nluc) 24 h after αRep expression induction with doxycycline. Virus replication was quantified by measurement of the Nluc enzymatic activity. Data are mean± s.e.m. n=2 independent infections with n=3 technical replicates.

### αReps C3 and F3 inhibit FluPol transcription

As shown above, αReps C3 and F3 inhibit the FluPol transcription + replication activity when expressed intracellularly (**Fig. 4B**). To determine if C3 and F3 inhibit more specifically transcription, we assessed in the same assay the transcriptional-only activity of a FluPol mutant with a single mutation in PA (PA-C95A) that is defective for replication (Nilsson-Payant et al., 2018). **Fig. 7** showed that FluPol transcription is inhibited by both αReps, C3 being more effective than F3 when compared with their activities against the FluPol wild-type form, suggesting that C3 may affect more specifically transcription than its anti-627 domain homolog.

**Figure 7:**
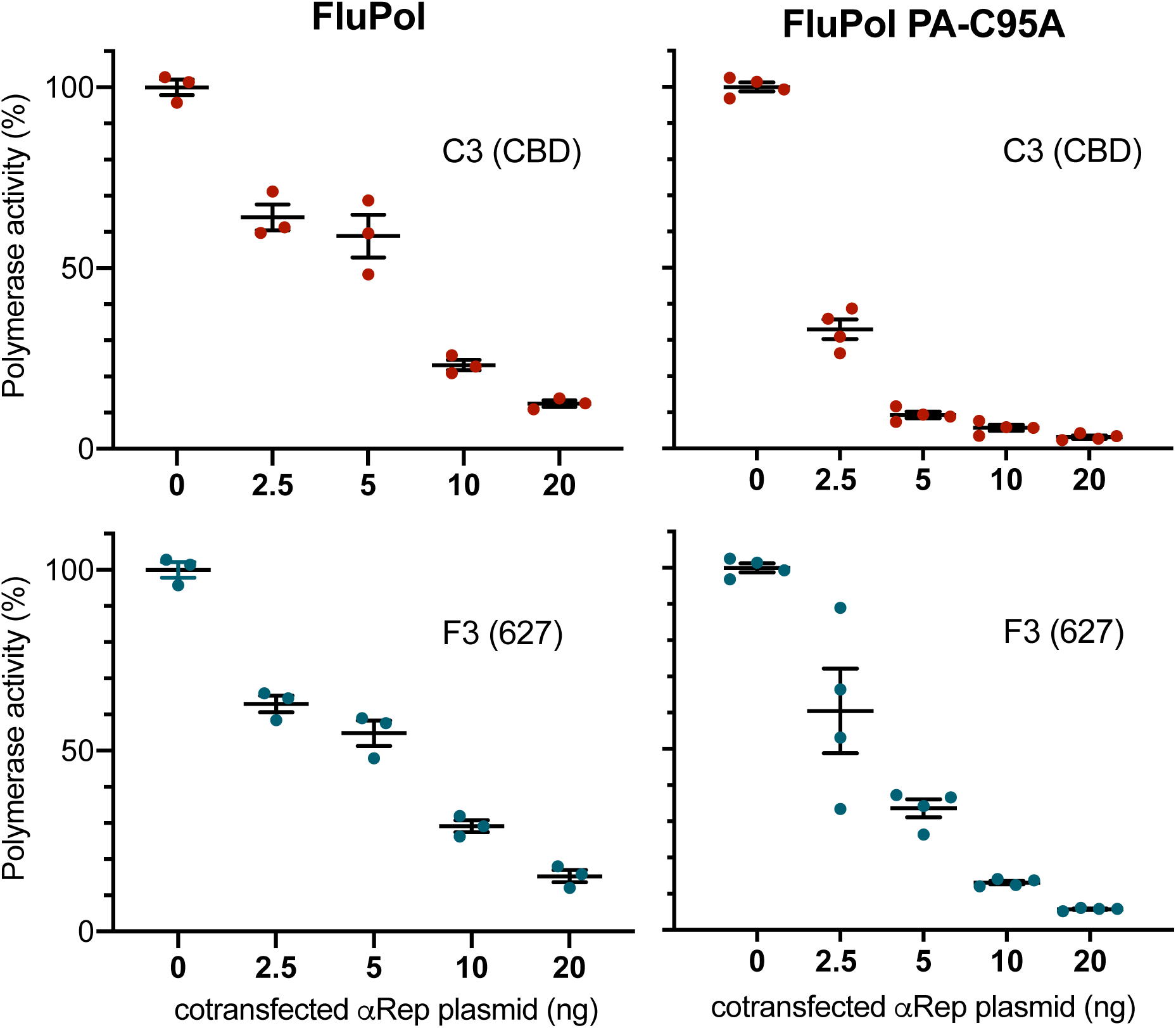
Effect of αRep C3 and F3 expression on the FluPol replication and/or transcription activities. Using a luciferase-reporter minireplicon assay, the FluPol activity was measured in transfection containing 0 to 50 ng of the C3 (CBD) or the F3 (627) plasmids per P96 well. An additional empty plasmid was included in each transfection to transfect the same amount of DNA. Left panels: RNA-polymerase activity was measured with a wild-type form of FluPol (WSN strain). Right panels: RNA-polymerase activity was measured with the mutated form of FluPol (FluPol PA-C95A in which PA was substituted by PA C95A) which is deficient in replicase activity (Nilsson-Payant et al., 2018). Data are mean± s.e.m. with n=3 or 4 technical replicates.

### αReps C3 and F3 do not interfere with PB2 targeting to the nucleus

The inhibition of the RNA-polymerase activity of FluPol by the αReps could result from a block in the genome transcription/replication process or in any upstream step, such as the assembly of the polymerase subunits or the import of the polymerase subunits in the nucleus. To determine if the two αReps may interfere with the translocation of FluPol to the nucleus, we first checked the cellular localization of C3 and F3 by confocal microscopy. We transfected the eukaryotic expression plasmids driving their expression in HEK293T cells. C3 and F3 αReps were mainly found in the cytoplasmic compartment (**S3 Fig**). Next, we determined whether C3 and/or F3 may interfere with the nuclear import of the PB2 subunit into the nucleus. We thus used plasmids to co-express a Flag-tagged PB2 with HA-tagged αReps to visualize the localization of the PB2 subunit in presence of these two binders. Our experiments showed that PB2 remained predominantly nuclear when C3 or F3 were co-expressed (**Fig. 8**). No retargeting of C3 and F3 αReps to the nucleus was evidenced when PB2 was co-expressed.

**Figure 8:**
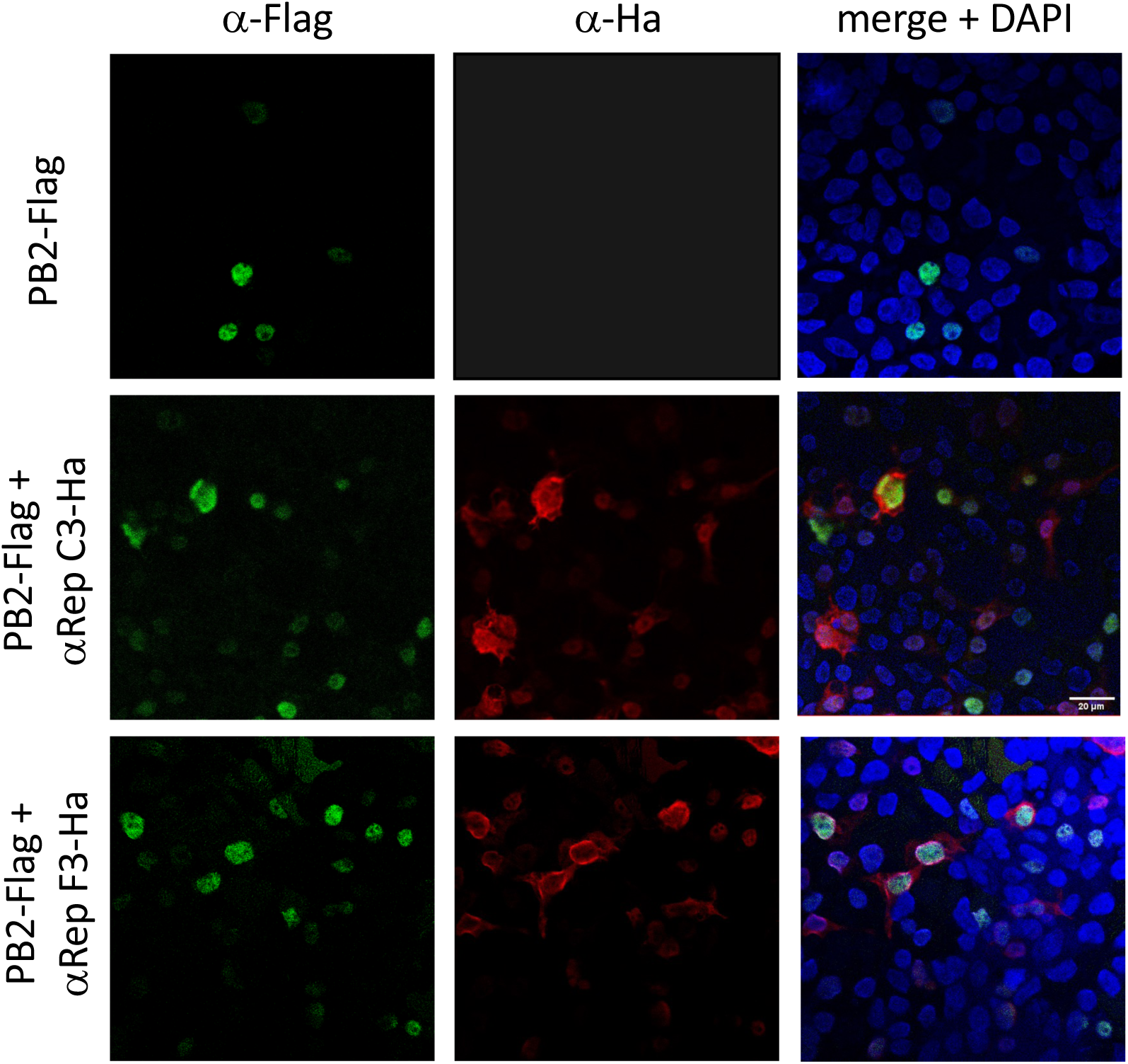
Subcellular localization of the PB2 subunit when coexpressed with αReps C3 and F3. PB2 was tagged with the epitope Flag and revealed with an anti-Flag antibody. αReps C3 and F3 were HA-tagged and revealed with an anti-HA antibody.

### C3-F3 dimerization increases their intrinsic activity

In order to determine if multimerization may promote synergy in their neutralization activity, we aimed at generating multivalent αReps. To build the C3-F3 and F3-C3 heterodimers, we inserted a 25-amino acid long linker (GGGGS)_5_ between these two subunits. This linker length (that can reach 8 nm in length) allows the binding of these two heterodimers on the monomeric forms of FluPol and on the replicative-competent dimeric form of FluPol (Carrique et al., 2020; Fan et al., 2019). Interestingly, a four-fold synergic effect in the inhibition efficiency on the FluPol activity was evidenced when C3 and F3 were covalently linked in both C3-F3 and F3-C3 constructs (**Fig. 4B**). The two chimera (C3-F3 and F3-C3) exhibited a slightly higher neutralization activity than their monomeric counterparts (**Fig. 4C)**. Thus, linkage of C3 and F3 markedly increases their FluPol inhibitory potency.

## Discussion

To date, only intracellular single domain antibodies (also known as intrabodies or VHHs) were screened as nanobinders to discover sites on the surface of FluPol that are sensitive to inhibition (Keown et al., 2022; Bessonne et al., 2024). Thus, following immunization of a llama and phage display screening, two VHHs (Nb8191 and VHH16) isolated in these two studies were found to bind a critical site for FluPol functioning (defined by the N-ter of PB1 and an adjacent PA α-helix). αReps offer another type of high-affinity binders that can be selected by *in vitro* techniques (phage display) without immunization. αReps are engineered artificial proteins, whose scaffold is derived from naturally occurring ankyrin repeats. A main advantage of αReps is their solubility and stability even in reducing conditions such as in the intracellular environment where they could block different steps of the virus cycle. They can either target viral glycoproteins, such as the spike of SARS-CoV-2 to block infection (Thébault et al., 2022) or bind to intracellular viral proteins like the HIV-1 nucleocapsid and to negatively interfere with virus maturation (Hadpech et al., 2017).

As an alternative to VHH, we screened an αReps library to identify additional sensitive targets on the FluPol surface. The structurally well-defined modules of the RNA-polymerase of influenza viruses (its central core constituted by PB1 subunit and PA-C and the flexibly-linked PB2 domains, CBD, 627 and NLS) were thus used in four independent screening. While we did not evidence anti-core and anti-NLS domain αReps able to interfere with the FluPol functioning, a high proportion of αReps specific of the 627-domain displays efficient inhibitory activity. One of them (αRep F3) displayed with the αRep C3 specific of the CBD high FluPol inhibitory and virus neutralizing activities when expressed in the cytosol. We also demonstrated the simplicity of αRep bioengineering to increase their inhibitory activity using multivalent forms. The C3-F3 and F3-C3 dimers were more effective than their monomeric counterparts to inhibit FluPol functioning and neutralize influenza virus. We also explored the ability of the two αReps to block virus replication when their expression was induced at different times post-infection. Induction at a late time in the virus cycle (6 hrs) resulted in a reduction of 40-50% of virus production, suggesting that the block may occur at late stages of the virus cycle. What could be the mode of action of αReps C3 and F3 to neutralize intracellularly influenza viruses? While we did not obtain clear evidence of a block of the transport of PB2 to the nucleus in the cells expressing the αReps, we found that they were able to block transcription, suggesting that they may prevent FluPol from making interactions in the nucleus with critical viral or host factors or adopting functional conformations. Considering the high structural versatility of FluPol in its functioning (Pflug et al., 2014; Reich et al., 2014; Thierry et al., 2016), we favor this last hypothesis. However, the relative high number of anti-627 αReps able to block FluPol polymerase activity may also suggest several modes of action, some of them being able to block interaction with nuclear proteins while others may block FluPol structural transitions. Structural characterization of the complexes between FluPol and inhibitory αReps should provide key information of the *modus operandi* of these ligands to block virus replication.

C3 and F3 resulted in efficient neutralization of viruses belonging to the different lineages (human H1N1 and H3N2 viruses and an avian H7N1 virus), a feature that may result from the high sequence conservation on FluPol sequences among influenza A viruses, and possibly on their high affinity for their targets. C3 and F3 binding sites may represent options for designing new types of influenza antivirals.

To conclude, we selected biosynthetic proteins (αReps) as specific and versatile neutralizing binders targeting different mobile domains of FluPol when expressed intracellularly. These biosynthetic proteins provide starting points for the identification of target sites to define new strategies to design influenza antivirals able to mimic anti-FluPol αReps activities. To our knowledge, it is the first time that the 627-domain has been identified as targetable to design antivirals. Our data also suggest that using bi-specific binders should also be considered to increase their activities and overcome emergence of escape variants.

## Materials and methods

### Cells

Human embryonic kidney 293T (HEK293T) were maintained in Dulbecco’s modified Eagle medium (DMEM) + 10% Fetal Calf Serum. High Five insect cells were grown in Express Five media (Life Technologies). Rabbit kidney (RK13) cells and derived cell clones were grown in modified Eagle’s medium (MEM, Eurobio scientific) supplemented with 5% fetal calf serum, glutamine, and penicillin-streptomycin. For RK13-derived clones, 1 μg/ml of puromycin was added to the cell culture medium.

### FluPol PA-PB1 heterodimer and PB2 domains expression and purification

Expression of the FluPol heterodimer PA(197-716) and PB1(1-696) has been previously described (Swale et al., 2016). Briefly, large scale High Five insect cells suspension cultures were infected at 0.2% (V/V) with the mother solution of a recombinant baculovirus governing expression of the fusion protein TEV - PA(197-716) - PB1(1-686) - GFP with TEV cleavage sites located between TEV and PA, PA and PB1 and PB1 and GFP. A His-tag was positioned in front of PA. Cultures were maintained at 0.5 – 1 × 10^6^ cells/mL until proliferation arrest (24–48 h after infection). To recover the polyprotein derived-products, cells were spun down at 800 g for 10 min and pellets were stored at − 80 °C. The PA-PB1 heterodimer was purified using the protocol described in Swale et al., 2016.

His-tagged PB2 C-terminal separate domains (**S1 Fig**) were produced in Rosetta (D3) bacteria. Briefly, bacterial clones containing the plasmids encoding the PB2 domains were cultured in large scale suspensions under agitation at 37°C until the optical density reached 0.6. Next, Rosetta cells were maintained under gentle agitation overnight at 17°C in the presence of 0.5 mM IPTG. Bacteria were then pelleted by centrifugation (5.000 x g for 30 min at 4°C) and bacterial cell pellets were resuspended in 50mM Tris, 500mM NaCl, pH 7.4 to 8, containing a cocktail of protease inhibitors (Roche Diagnostics GmbH). Bacteria maintained at 4°C were lysed by sonication (5 times x (30 s sonication at around 40% sonication amplitude and 30 s rest)) using a Q700 Sonicator (QSONICA). Bacterial cells lysates were clarified by centrifugation at 10,000 × g for 30 min at 4°C. Soluble PB2 C-terminal domains were purified by affinity chromatography on HisTrap columns (GE Healthcare Life Sciences) and analyzed by SDS-PAGE. Fractions of interest were pooled and injected in a Gel filtration Superdex S200 previously equilibrated with 20mM Tris, 500mM NaCl buffer. Fractions containing purified PB2 C-domains were pooled and frozen at −80°C.

### Screening of the αRep library against the PA-PB1 dimer and PB2 domains

The construction of the αRep phage library 2.1 has been previously described (Guellouz et al., 2013). The αRep library was constructed by polymerization of synthetic microgenes corresponding to individual HEAT-like repeats, and the αRep proteins were expressed at the surface of M13-derived filamentous phages. The library is estimated to contain 1.7 x 10^9^ independent clones. The αRep library screening was carried out as described (Hadpech et al., 2017) with minor modifications. Purified PA-PB1 dimer and PB2 domains diluted at 1 mM in PBS containing 0.05% Tween-20 (PBST) was immobilized on Ni++-NTA-biotin streptavidin-coated 96-well ELISA plate by incubation overnight at 4°C in a moisture chamber. The coated wells were washed four times with PBST, and saturated with blocking solution (2% BSA in PBST; 200 µL/well) for 1h, after which an aliquot of the phage library was added to the wells coated with either the PA-PB1 heterodimer or the PB2 domains, and incubated at room temperature for 2h with shaking. Next, wells were extensively washed in PBST, and bound phages were eluted by three successive rounds of adsorption/elution. Phage elution was performed by an acidic glycine solution (0.1 M glycine-HCl, pH 2.5) and buffered using Tris 1 M. The population of αRep-displayed phages eluted from the FluPol baits was amplified and subcloned in a XL-1 Blue cells. Individual phage clones were selected and amplified as previously described (Guellouz et al., 2013, Valerio-Lepiniec et al., 2015), and their respective binding activity towards the protein targets was determined by ELISA. 100 μL-aliquots of purified targets were diluted in PBS and loaded into the wells of a Ni^2+^-NTA-biotin streptavidin-coated ELISA plate, then incubated overnight at 4°C. The coated plate was washed four times with TBST, then blocked with PBST-BSA (200 μL per well) for 1 hour with shaking. After a washing step, 100 μL-aliquots of each phage culture supernatant were added to the wells and incubated at room temperature for 1 hour, followed by HRP-conjugated mouse anti-M13 (Interchim) diluted to 1:2,000 in TBST-BSA (100 μL-aliquot per well), and incubation proceeded for an extra 1 hour. The wells were washed again, prior to the addition of 100 μL BM Blue POD soluble Substrate (Roche). Reaction was stopped with 1 N HCl, and absorbance measured at 450 nm. Phage clones showing a high binding activity towards the immobilized target were sequenced and kept for cytoplasmic expression of individual αRep proteins.

### αReps expression in eukaryotic cells

#### Transient expression

The αRep genes encoding the FluPol binders were subcloned in a modified version of the eukaryotic expression vector pcDNA3 in which its multiple cloning site was modified to insert a HA-tag encoding sequence at the 3’-end and allowing the in frame cloning of αRep encoding sequences at BamHI and HindIII restriction sites. ***Inducible expression in stable cell clones.*** RK13 cell clones expressing HA-tagged C3 or F3 αReps in a doxycycline-inducible manner were generated by co-transfection of pCMV-TET3G, pTRE3G-αRep and a plasmid expressing the puromycin resistance cassette and selected in the presence of puromycin (1 μg/mL).

### Minireplicon assay

293T cells were seeded in P96 wells (7 x 10^4^ cells/well) and transfected on day 1 with plasmids driving the expression of the viral proteins PB1, PB2, PA NP, and pPolI-WSN-NAfirefly luciferase with a pcDNA3 plasmid derivate encoding an HA-tagged αRep as previously described (Leymarie et al., 2013). Plasmid pCMV-Nanoluc (Promega) was used as an internal control for transfection efficiency. As a negative control, 293T cells were transfected with the same plasmids, with the exception of the PA expression plasmid. After transfection, the cells were incubated at 37°C for 48 h, and then luciferase activity was measured with the Luciferase Assay System (Promega) on a Tecan Infinite M200Pro luminometer according to the manufacturer’s instructions.

### Reporter influenza and vesicular stomatitis viruses

The recombinant reporter virus WSN(H1N1)-nanoluciferase (H1N1-Nluc, previously named PASTN or PA-SWAP-2A-NLuc50 in (Tran et al., 2013) was kindly provided by Andrew Mehle. In this reporter virus, the “self-cleaving” 2A peptide from porcine teschovirus and the nanoluciferase coding sequence were placed downstream of the PA sequence to create a contiguous ORF. Native packaging sequences were restored by repeating the terminal 50 nucleotides of the PA ORF (including the stop codon) after the Nluc stop codon adjacent the native untranslated region. Direct repeats were removed from the reporter gene by introducing silent mutations at the 3’ end of the PA ORF. Reverse genetic systems for avian influenza A/Turkey/Italy/977/1999 [H7N1] and human influenza A/Scotland/20/1974 [H3N2] viruses have been previously elaborated and used to generate the recombinant reporter viruses H7N1-Nluc and H3N2-Nluc (Mettier et al., 2021; Meyer et al., 2016). Both these Nluc reporter viruses were designed using the same strategy than H1N1-Nluc, with their PA segments encoding a PA-2A-Nluc polyprotein (Bessonne et al., 2024). The reporter VSV-mCherry virus (Heilmann et al., 2019) in which the fluorescent protein mCherry ORF was inserted in the L-protein reading frame was kindly provided by Emmanuel Heilmann.

### Immunofluorescence assay

HEK-293T cells were seeded in 24-well plates on coverslips one day before transfection. Transfections were carried out with Lipofectamine 2000 (Thermo Fisher Scientific) using 500ng of the pcDNA3-derived plasmid encoding the αRep fused to an HA-tag with or without the pcDNA3-derived encoding PB2 tagged with the Flag sequence. Twenty-four hours later, cells were fixed with 4% paraformaldehyde (PFA). αReps were revealed using a secondary anti-HA antibody coupled to Alexa Fluor 546 and PB2 with a secondary antibody coupled to Alexa Fluor 488. Coverslips were then visualized by confocal microscopy.

### Co-immunoprecipitation assay

HEK-293T cells were seeded in 6-well plates 24h before transfection. Transfections were carried out with mixtures of the following plasmids: pcDNA3-αRep-HA, pCI-PB2-V5, pCI-PA, pCI-PB1 and pCI (for a total of 2 μg/well). Cells were harvested 24 h post-transfection and lysed with a buffer containing 50 mM Tris-HCl pH7.4, 150 mM NaCl, 0.5% NP40 and a cocktail of protease inhibitors (#11-836-170-001 Roche Diagnostic). Samples were incubated for 30 min on ice and cleared by centrifugation at 16,000 x g for 10 min at 4°C). Supernatants were subjected to immunoprecipitation using an anti-HA antibody at 4°C for 1 h before addition of magnetic protein A beads (Pierce) previously equilibrated with the lysis buffer. Magnetic beads were washed three times with the same buffer. The beads were resuspended in Laemmli’s buffer prior to Western blot analysis.

### Western Blot assay

Immunoprecipitates were loaded on a 12% polyacrylamide gel for electrophoresis. Material present in the gel was transferred onto a nitrocellulose membrane. The membrane was blocked with 5% skimmed milk in Phosphate buffered saline (PBS) for 30 min and incubated overnight at 4°C with antibodies. Membranes were then washed extensively in PBS-0.1% Tween 20 and incubated for 1 h with HRP-conjugated IgG secondary antibodies at room temperature (RT). After washing, the membranes were scanned with an Chemidoc imaging system (Biorad).

## Acknowledgments

M.B. acknowledges fellowships of the DIM1Health and of the Animal Health Division of INRAE, J.M. acknowledges fellowships of the ANR program and the Animal Health Division of INRAE. B.D. acknowledges support of the ANR-17-CE18-0006-01 program. We thank Andrew Mehle for the gift of the reverse genetics system of the IAV H1N1-Nluc, Ronan Le Goffic for the gift of the H7N1-Nluc and the H3N2-Nluc viruses. We thank Nicolas Meunier for his help in statistical analyses. This work has benefited from the ALPHAREP Platform of I2BC, supported by French Infrastructure for Integrated Structural Biology (ANR-10-INBS-05). The present work has benefited from Imagerie-Gif core facility supported by the Agence Nationale de la Recherche (ANR-10-INBS-04/FranceBioImaging ; ANR-11-IDEX-0003-02/ Saclay Plant Sciences).

## Supplementary figure legends

**S1 Figure:**
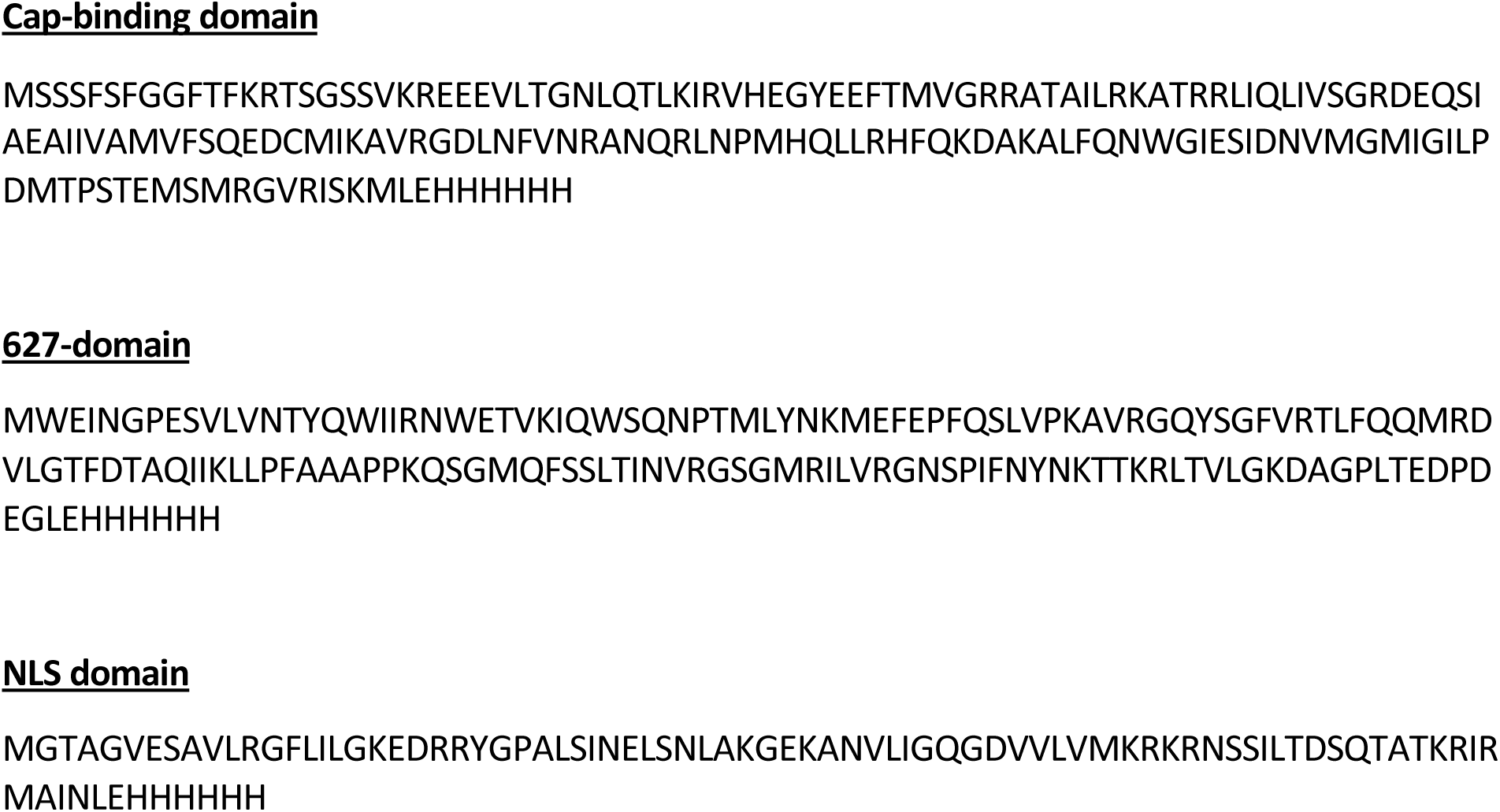
Amino acid sequences of the PB2 domains used for αReps screenings.

**S2 Figure:**
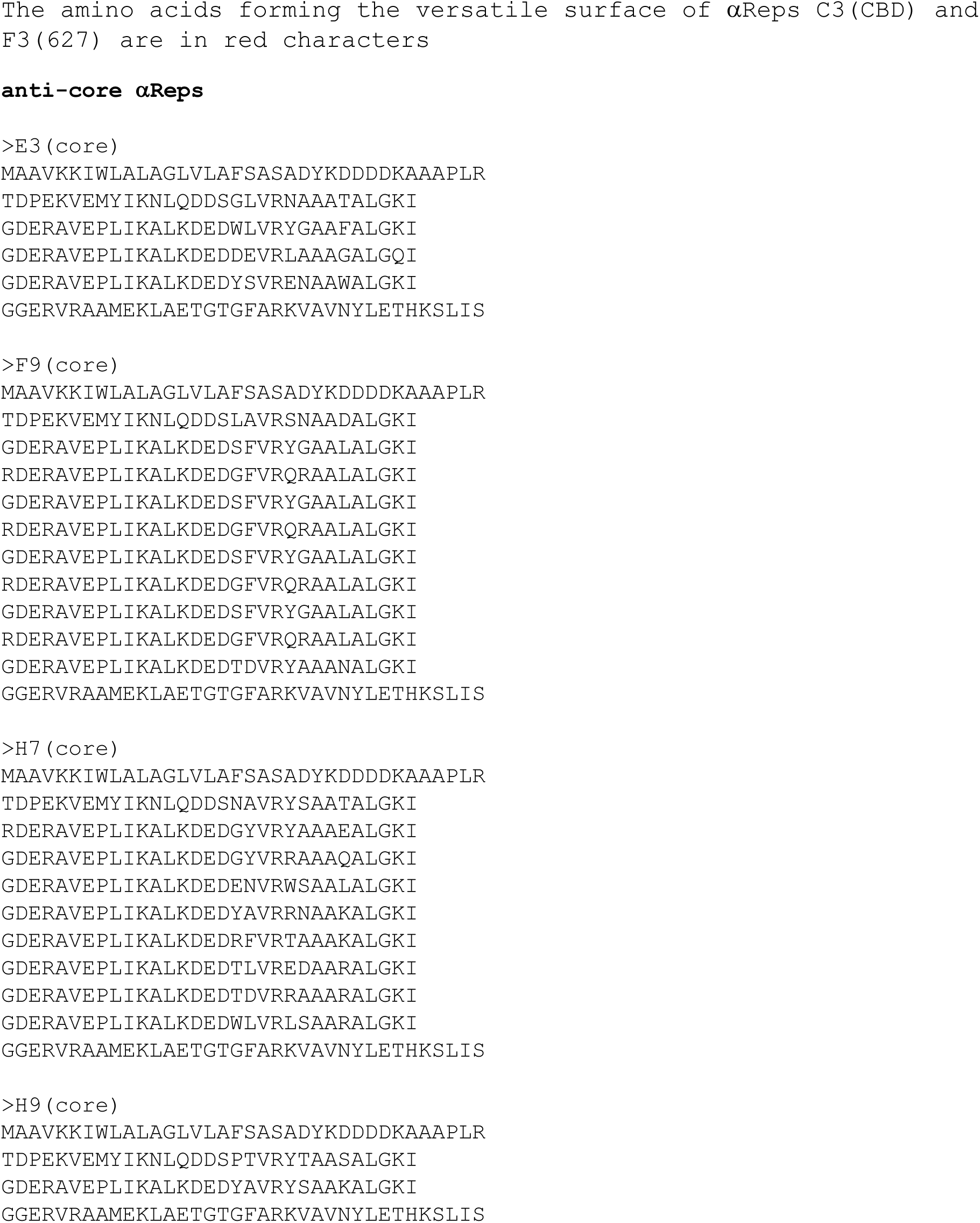
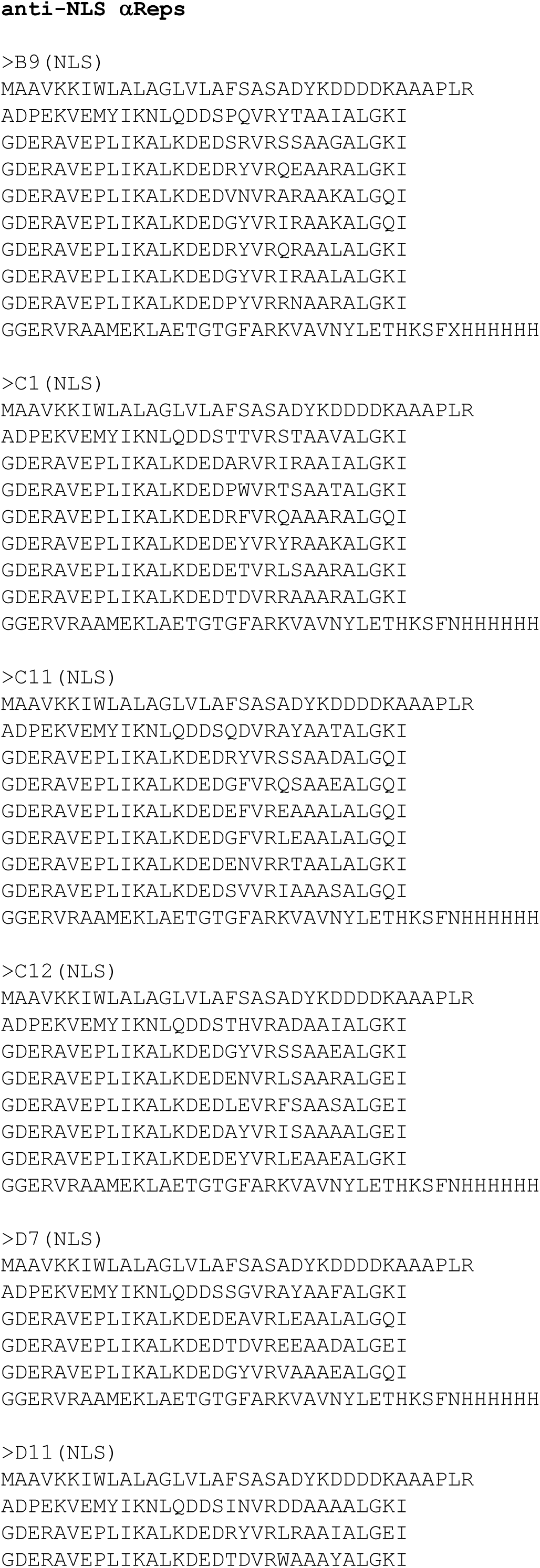
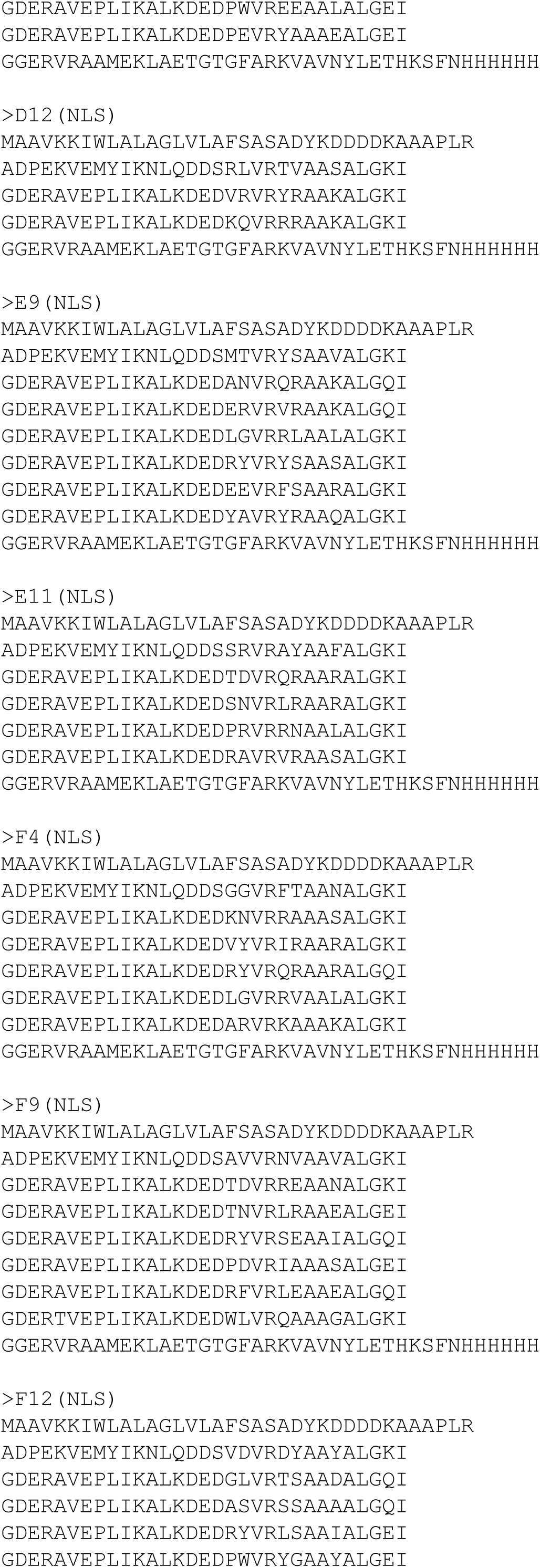
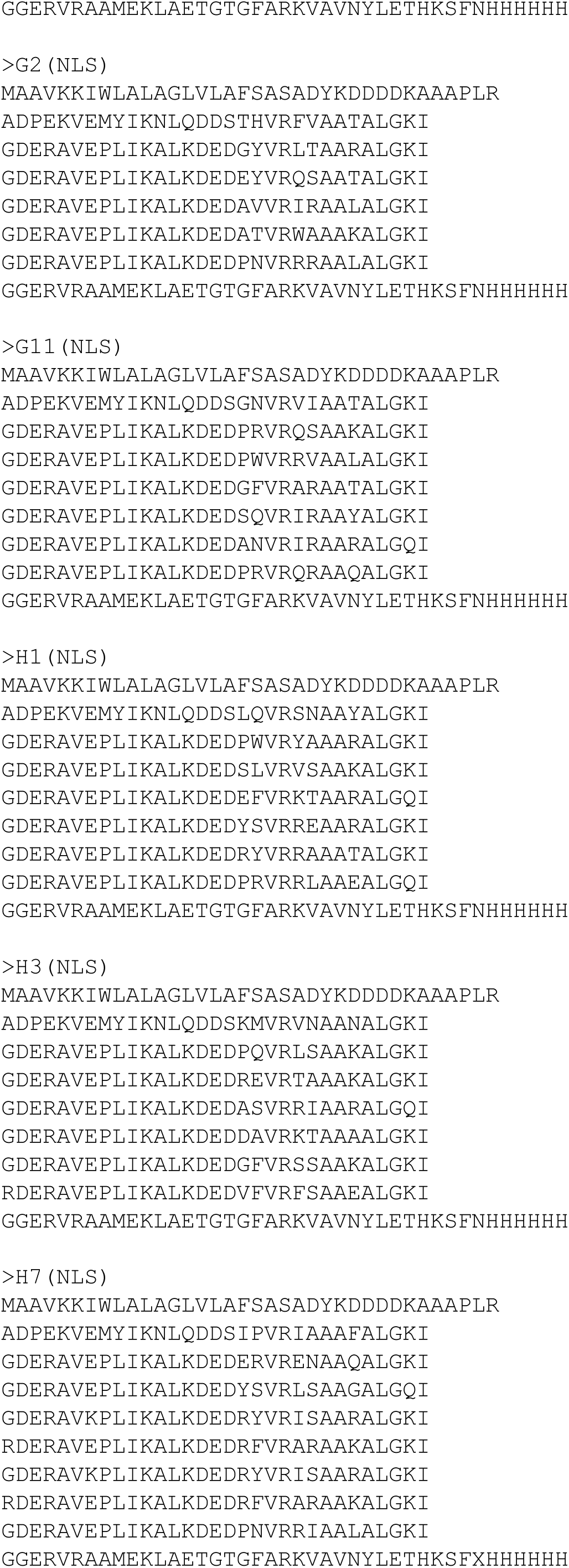
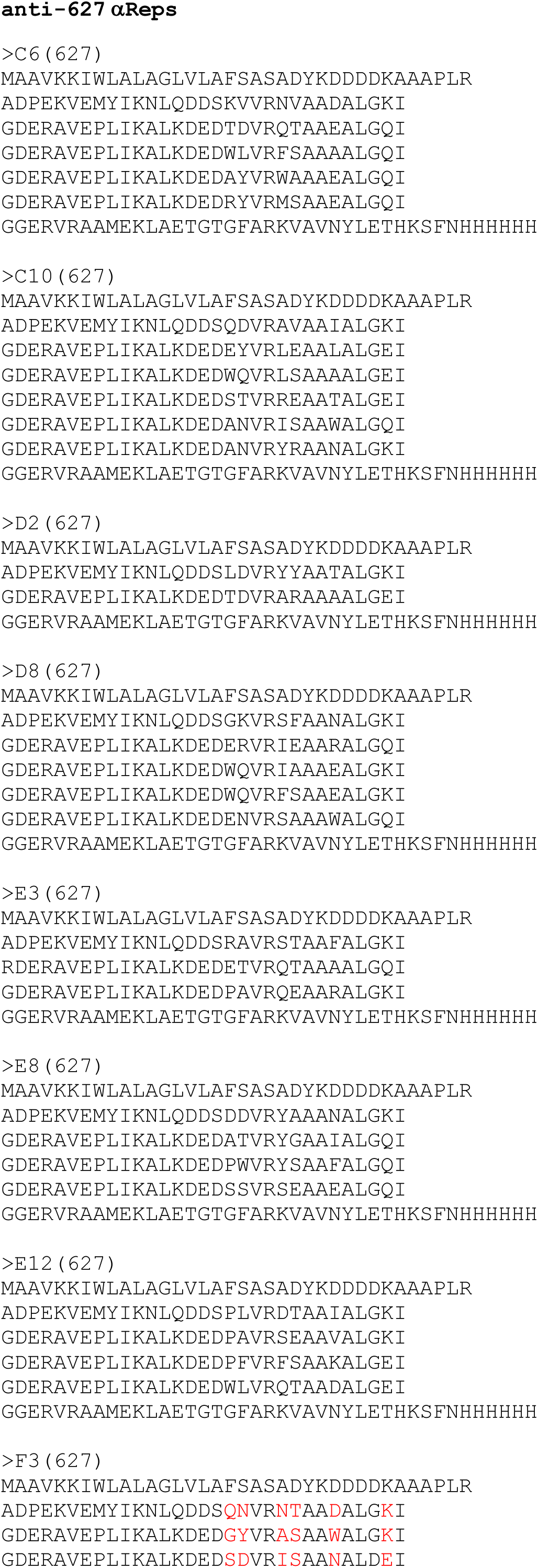
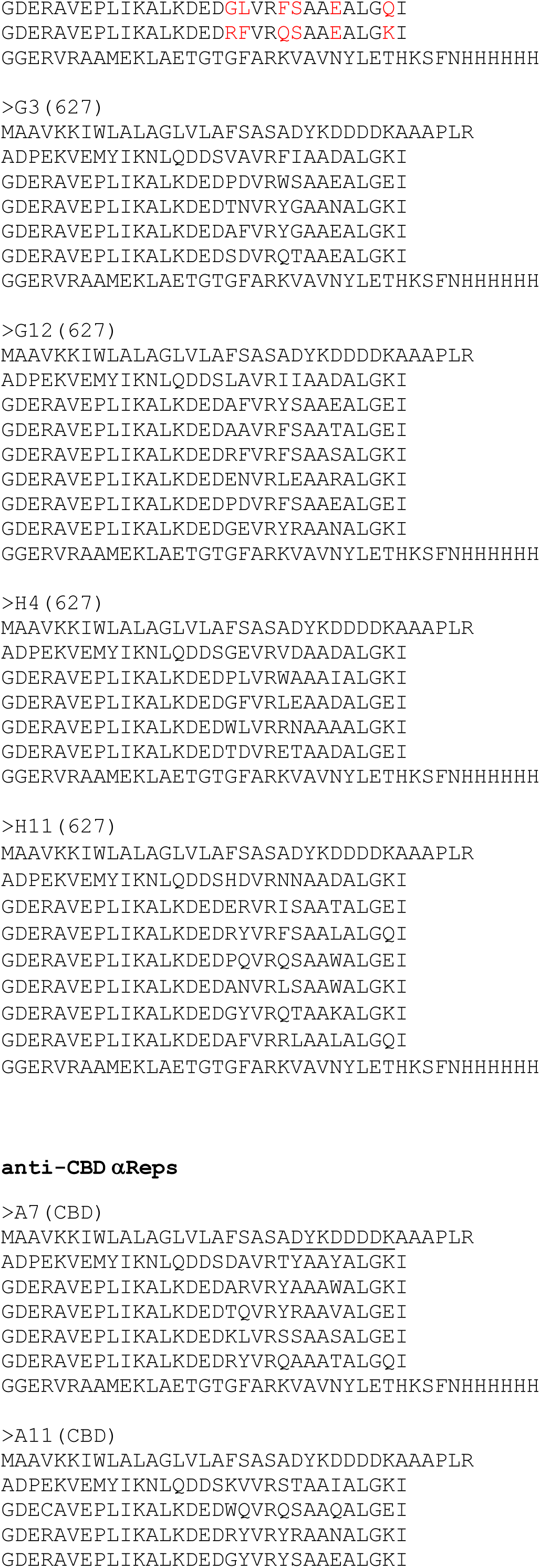
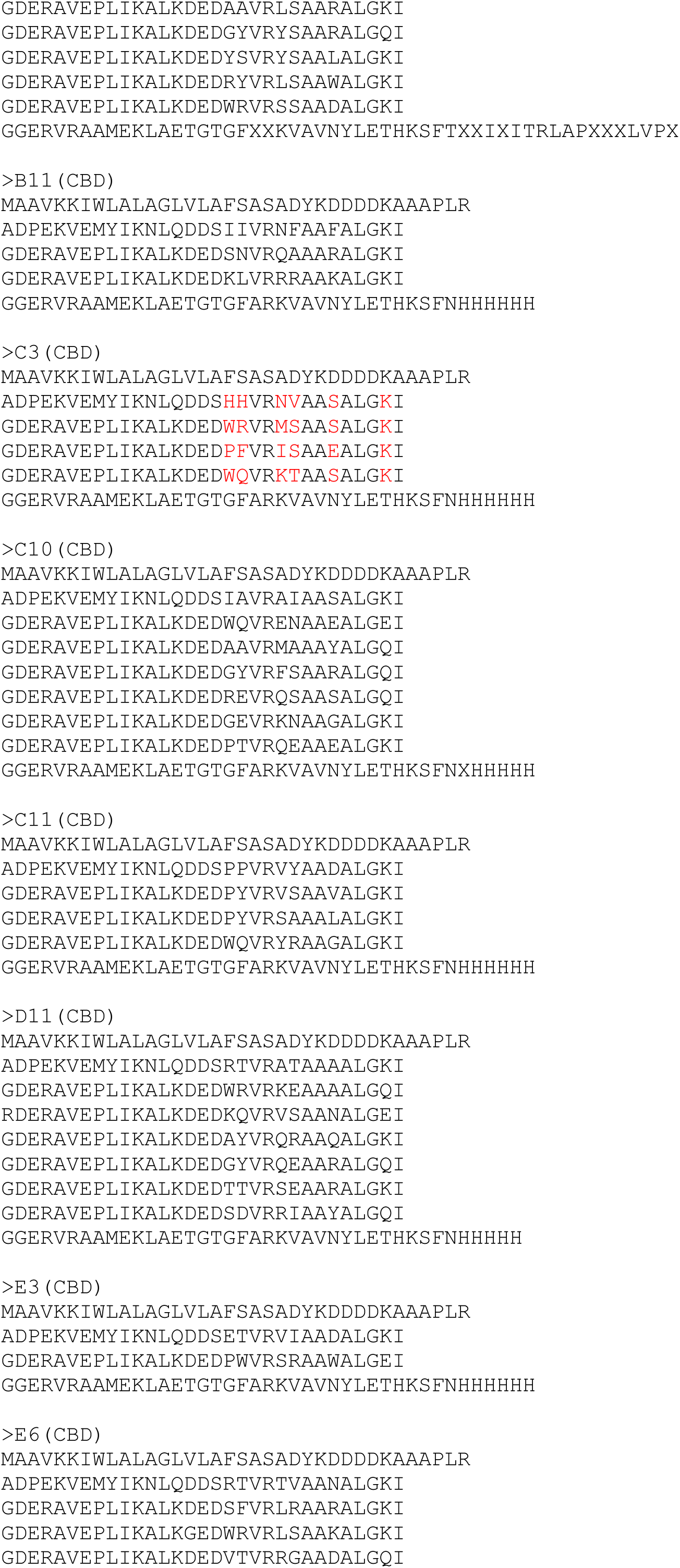
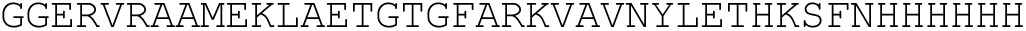
Specificity and αReps amino acid sequences.

**S3 Figure:**
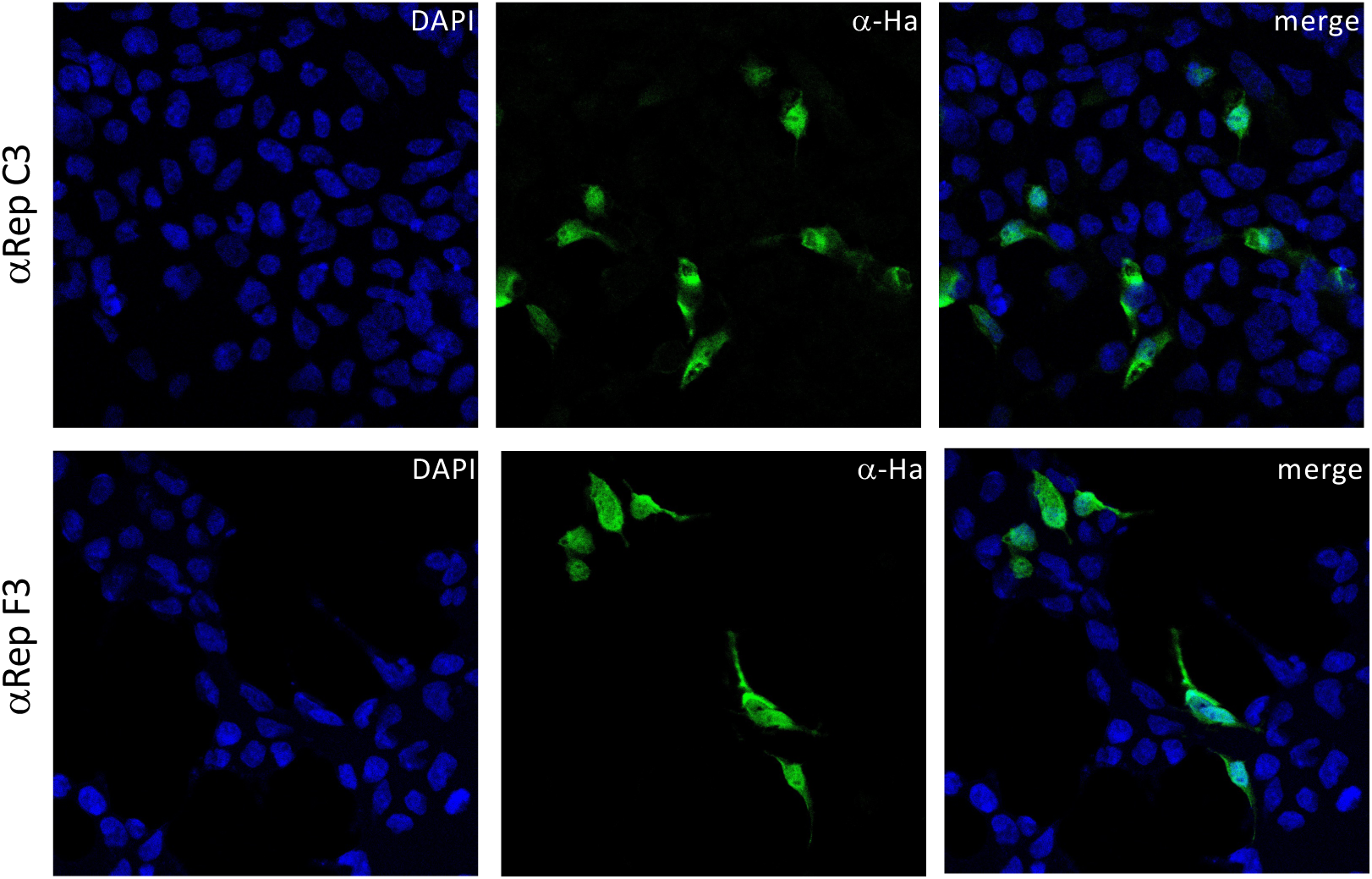
Indirect immunofluorescence assay to reveal HA-tagged αReps.

**S4 Figure:**
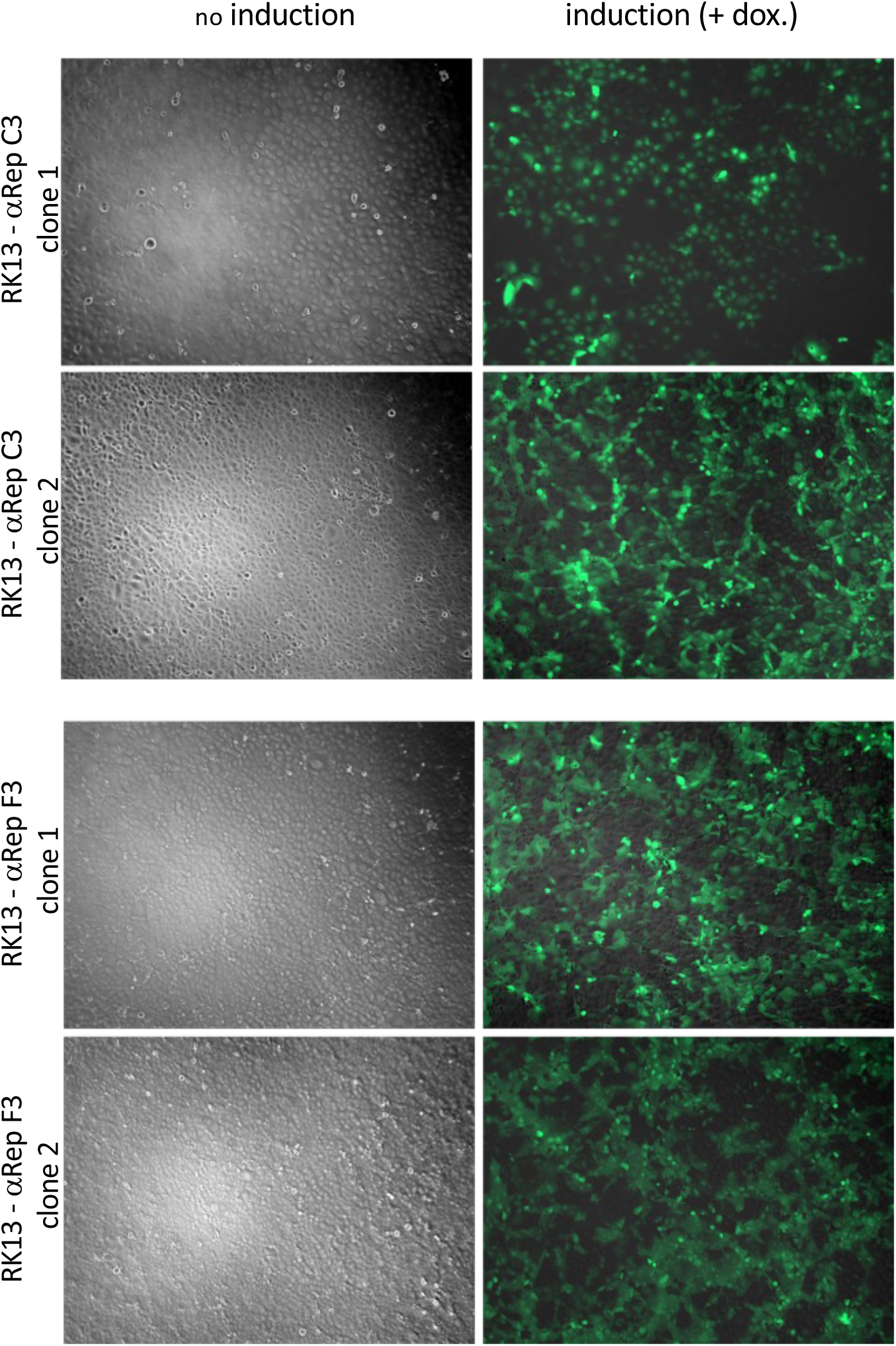
Inducible αReps expression after addition of doxycycline in two RK13 clones for αReps C3 and F3.

## References

Andrade, M.A., Petosa, C., O’Donoghue, S.I., Müller, C.W., Bork, P., 2001. Comparison of ARM and HEAT protein repeats11Edited by P. E. Wright. J. Mol. Biol. 309, 1–18. 10.1006/jmbi.2001.4624

Bessonne, M., Morel, J., Nevers, Q., Ballandras-Colas, A., Grange, M., Roussel, A., Crépin, T., Delmas, B., 2023. Antiviral activity of intracellular nanobodies targeting the influenza virus RNA-polymerase core. PLOS Pathog. 20, e1011642. 10.1371/journal.ppat.1011642

Binz, H.K., Amstutz, P., Plückthun, A., 2005. Engineering novel binding proteins from nonimmunoglobulin domains. Nat. Biotechnol. 23, 1257–1268. 10.1038/nbt1127

Carrique, L., Fan, H., Walker, A.P., Keown, J.R., Sharps, J., Staller, E., Barclay, W.S., Fodor, E., Grimes, J.M., 2020. Host ANP32A mediates the assembly of the influenza virus replicase. Nature 587, 638–643. 10.1038/s41586-020-2927-z

Chevrel, A., Urvoas, A., de la Sierra-Gallay, I.L., Aumont-Nicaise, M., Moutel, S., Desmadril, M., Perez, F., Gautreau, A., van Tilbeurgh, H., Minard, P., Valerio-Lepiniec, M., 2015. Specific GFP-binding artificial proteins (αRep): a new tool for in vitro to live cell applications. Biosci. Rep. 35, e00223. 10.1042/BSR20150080

Da Costa, B., Sausset, A., Munier, S., Ghounaris, A., Naffakh, N., Le Goffic, R., Delmas, B., 2015. Temperature-Sensitive Mutants in the Influenza A Virus RNA Polymerase: Alterations in the PA Linker Reduce Nuclear Targeting of the PB1-PA Dimer and Result in Viral Attenuation. J. Virol. 89, 6376–6390. 10.1128/JVI.00589-15

Deng, T., Sharps, J., Fodor, E., Brownlee, G.G., 2005. In Vitro Assembly of PB2 with a PB1-PA Dimer Supports a New Model of Assembly of Influenza A Virus Polymerase Subunits into a Functional Trimeric Complex. J. Virol. 79, 8669–8674. 10.1128/jvi.79.13.8669-8674.2005

Fan, H., Walker, A.P., Carrique, L., Keown, J.R., Serna Martin, I., Karia, D., Sharps, J., Hengrung, N., Pardon, E., Steyaert, J., Grimes, J.M., Fodor, E., 2019. Structures of influenza A virus RNA polymerase offer insight into viral genome replication. Nature 573, 287–290. 10.1038/s41586-019-1530-7

Guellouz, A., Valerio-Lepiniec, M., Urvoas, A., Chevrel, A., Graille, M., Fourati-Kammoun, Z., Desmadril, M., van Tilbeurgh, H., Minard, P., 2013. Selection of Specific Protein Binders for Pre-Defined Targets from an Optimized Library of Artificial Helicoidal Repeat Proteins (alphaRep). PLoS ONE 8, e71512. 10.1371/journal.pone.0071512

Hadpech, S., Nangola, S., Chupradit, K., Fanhchaksai, K., Furnon, W., Urvoas, A., Valerio-Lepiniec, M., Minard, P., Boulanger, P., Hong, S.-S., Tayapiwatana, C., 2017. Alpha-helicoidal HEAT-like Repeat Proteins (αRep) Selected as Interactors of HIV-1 Nucleocapsid Negatively Interfere with Viral Genome Packaging and Virus Maturation. Sci. Rep. 7, 16335. 10.1038/s41598-017-16451-w

Heilmann, E., Kimpel, J., Geley, S., Naschberger, A., Urbiola, C., Nolden, T., von Laer, D., Wollmann, G., 2019. The Methyltransferase Region of Vesicular Stomatitis Virus L Polymerase Is a Target Site for Functional Intramolecular Insertion. Viruses 11, 989. 10.3390/v11110989

Huet, S., Avilov, S.V., Ferbitz, L., Daigle, N., Cusack, S., Ellenberg, J., 2010. Nuclear Import and Assembly of Influenza A Virus RNA Polymerase Studied in Live Cells by Fluorescence Cross-Correlation Spectroscopy. J. Virol. 84, 1254–1264. 10.1128/jvi.01533-09

Keown, J.R., Zhu, Z., Carrique, L., Fan, H., Walker, A.P., Serna Martin, I., Pardon, E., Steyaert, J., Fodor, E., Grimes, J.M., 2022. Mapping inhibitory sites on the RNA polymerase of the 1918 pandemic influenza virus using nanobodies. Nat. Commun. 13, 251. 10.1038/s41467-021-27950-w

Leymarie, O., Jouvion, G., Hervé, P.-L., Chevalier, C., Lorin, V., Lecardonnel, J., Costa, B.D., Delmas, B., Escriou, N., Goffic, R.L., 2013. Kinetic Characterization of PB1-F2-Mediated Immunopathology during Highly Pathogenic Avian H5N1 Influenza Virus Infection. PLOS ONE 8, e57894. 10.1371/journal.pone.0057894

Mettier, J., Marc, D., Sedano, L., Da Costa, B., Chevalier, C., Le Goffic, R., 2021. Study of the host specificity of PB1-F2-associated virulence. Virulence 12, 1647–1660. 10.1080/21505594.2021.1933848

Meyer, L., Sausset, A., Sedano, L., Da Costa, B., Le Goffic, R., Delmas, B., 2016. Codon Deletions in the Influenza A Virus PA Gene Generate Temperature-Sensitive Viruses. J. Virol. 90, 3684–3693. 10.1128/jvi.03101-15

Nilsson-Payant, B.E., Sharps, J., Hengrung, N., Fodor, E., 2018. The Surface-Exposed PA51-72-Loop of the Influenza A Virus Polymerase Is Required for Viral Genome Replication. J. Virol. 92, e00687–18. 10.1128/JVI.00687-18

Pflug, A., Guilligay, D., Reich, S., Cusack, S., 2014. Structure of influenza A polymerase bound to the viral RNA promoter. Nature 516, 355–360. 10.1038/nature14008

Pflug, A., Lukarska, M., Resa-Infante, P., Reich, S., Cusack, S., 2017. Structural insights into RNA synthesis by the influenza virus transcription-replication machine. Virus Res. 234, 103–117. 10.1016/j.virusres.2017.01.013

Reich, S., Guilligay, D., Pflug, A., Malet, H., Berger, I., Crépin, T., Hart, D., Lunardi, T., Nanao, M., Ruigrok, R.W.H., Cusack, S., 2014. Structural insight into cap-snatching and RNA synthesis by influenza polymerase. Nature 516, 361–366. 10.1038/nature14009

Soetens, E., Ballegeer, M., Saelens, X., 2020. An Inside Job: Applications of Intracellular Single Domain Antibodies. Biomolecules 10, 1663. 10.3390/biom10121663

Swale, C., Monod, A., Tengo, L., Labaronne, A., Garzoni, F., Bourhis, J.-M., Cusack, S., Schoehn, G., Berger, I., Ruigrok, R.W., Crépin, T., 2016. Structural characterization of recombinant IAV polymerase reveals a stable complex between viral PA-PB1 heterodimer and host RanBP5. Sci. Rep. 6, 24727. 10.1038/srep24727

Thébault, S., Lejal, N., Dogliani, A., Donchet, A., Urvoas, A., Valerio-Lepiniec, M., Lavie, M., Baronti, C., Touret, F., Costa, B.D., Bourgon, C., Fraysse, A., Saint-Albin-Deliot, A., Morel, J., Klonjkowski, B., Lamballerie, X. de, Dubuisson, J., Roussel, A., Minard, P., Poder, S.L., Meunier, N., Delmas, B., 2022. Biosynthetic proteins targeting the SARS-CoV-2 spike as anti-virals. PLOS Pathog. 18, e1010799. 10.1371/journal.ppat.1010799

Thierry, E., Guilligay, D., Kosinski, J., Bock, T., Gaudon, S., Round, A., Pflug, A., Hengrung, N., El Omari, K., Baudin, F., Hart, D.J., Beck, M., Cusack, S., 2016. Influenza Polymerase Can Adopt an Alternative Configuration Involving a Radical Repacking of PB2 Domains. Mol. Cell 61, 125–137. 10.1016/j.molcel.2015.11.016

Tran, V., Moser, L.A., Poole, D.S., Mehle, A., 2013. Highly Sensitive Real-Time In Vivo Imaging of an Influenza Reporter Virus Reveals Dynamics of Replication and Spread. J. Virol. 87, 13321–13329. 10.1128/JVI.02381-13

Urvoas, A., Guellouz, A., Valerio-Lepiniec, M., Graille, M., Durand, D., Desravines, D.C., van Tilbeurgh, H., Desmadril, M., Minard, P., 2010. Design, Production and Molecular Structure of a New Family of Artificial Alpha-helicoidal Repeat Proteins (αRep) Based on Thermostable HEAT-like Repeats. J. Mol. Biol. 404, 307–327. 10.1016/j.jmb.2010.09.048

Valerio-Lepiniec, M., Urvoas, A., Chevrel, A., Guellouz, A., Ferrandez, Y., Mesneau, A., de la Sierra-Gallay, I.L., Aumont-Nicaise, M., Desmadril, M., van Tilbeurgh, H., Minard, P., 2015. The αRep artificial repeat protein scaffold: a new tool for crystallization and live cell applications. Biochem. Soc. Trans. 43, 819–824. 10.1042/BST20150075

Wandzik, J.M., Kouba, T., Karuppasamy, M., Pflug, A., Drncova, P., Provaznik, J., Azevedo, N., Cusack, S., 2020. A Structure-Based Model for the Complete Transcription Cycle of Influenza Polymerase. Cell 181, 877–893.e21. 10.1016/j.cell.2020.03.061

